# Polyether ionophore antibiotics target drug-resistant clinical isolates, persister cells, and biofilms

**DOI:** 10.1101/2023.02.13.528344

**Authors:** Malene Wollesen, Kasper Mikkelsen, Marie Selch Tvilum, Martin Vestergaard, Mikala Wang, Rikke L. Meyer, Hanne Ingmer, Thomas B. Poulsen, Thomas Tørring

**Affiliations:** Department of Chemistry, Aarhus University, Langelandsgade 140, DK-8000, Aarhus C, Denmark; Department of Biological and Chemical Engineering, Gustav Wieds Vej 10, DK-8000, Aarhus C, Denmark; Department of Veterinary and Animal Sciences, University of Copenhagen, Grønnegårdsvej 15, DK- 1870 Frederiksberg C, Denmark; Department of Clinical Microbiology, Aarhus University Hospital, Palle Juul-Jensens Boulevard 99, DK-8200 Aarhus N, Denmark; Interdisciplinary Nanoscience Center, Gustav Wieds Vej 14, DK-8000, Aarhus C, Denmark; Department of Biology, Ny Munkegade 114-116, DK-8000, Aarhus C, Denmark

**Keywords:** ionophore, antibiotics, biofilm, persister cells, antimicrobial resistance, staphylococcus aureus

## Abstract

Polyether ionophores are complex natural products known to transport various cations across biological membranes. While several members of this family are used in agriculture, e.g. as anti-coccidiostats, and have potent antibacterial activity, they are not currently pursued as antibiotics for human use. Polyether ionophores are typically grouped as having similar functions, despite the fact that they differ significantly in structure, and for this reason, it remains unclear how structure and activity are related. To triage whether certain members of the family constitute particularly interesting springboards for in-depth investigations, and future synthetic optimization, we here conduct a systematic comparative study of nine different polyether ionophores for their potential as antibiotics. This includes clinical isolates from bloodstream infections and studies of the compounds’ effects on bacterial biofilms and persister cells. We uncover distinct differences within the compound class and identify the compounds lasalocid, calcimycin, and nanchangmycin as having particularly interesting activity profiles for further development.

## Introduction

The number of infections caused by antimicrobial-resistant microorganisms is rising and poses a major threat to our society. A widely discussed survey from the antimicrobial resistance collaborators published in early 2022 estimated 4.97 million deaths globally associated with antimicrobial resistance in 2019 alone.^1^ The majority of currently used antibiotics can simplistically be said to target a limited number of processes common to most microorganisms: cell wall integrity (beta-lactams and glycopeptides), cell membrane (lipopeptides), translation (chloramphenicol and macrolides), transcription (rifamycins) and DNA replication (fluoroquinolones). Several papers have argued that new antimicrobial treatments may be discovered if we can alter the bacterial metabolic state as many antibiotics are only bactericidal to metabolically active bacteria.^2–4^ This is exemplified by the increase in bacterial uptake of aminoglycosides that is stimulated by the addition of simple metabolites such as fructose, mannitol, or glucose whose catabolism increases the proton motive force (PMF) and electric potential across the membrane (ΔΨ) and kill otherwise dormant bacteria.^5^ Similarly, it has been described that bicarbonate concentration differences between an infection and the typical *in vitro* screening in Mueller Hinton broth can affect drug translatability. Farha *et al*. elegantly showed that this is likely because bicarbonate dissipates the ΔpH component of the PMF causing a compensatory increase in the ΔΨ.^6,7^

These deviations from a simplistic model of an antibiotic blocking a single essential process begs a more nuanced view of the bacterial physiology during infections and the ultimate causes of bacterial death beyond the target-focused proximate causes as discussed by Baquero and Levin.^8^

The polyether ionophore natural products typically produced by Streptomycetes fit this particular description well. They are a large class of structurally diverse amphiphilic antibiotics with the ability to perturb cellular ion-gradients by effecting trans-membrane, electroneutral exchange of ions/protons down their concentration-gradients.^9–11^ The molecules achieve this by a dynamic conformational change upon binding the cation. This initial ion-complex then undergoes a conformational change to fully encapsulate the bound ion by orienting multiple polar groups toward the cavity interior and, conversely, hydrophobic groups to the exterior allowing diffusion of the ionophore-ion complex across the lipid bilayer.^12^ The action of ionophores on (bacterial) cells is dependent on multiple parameters that transcend this simple, canonical mechanism. Depending on the specific structure, ionophores have differential affinities for different cations and likely have differing ion-transport dynamics and membrane affinities. As such, exposure to a given ionophore will constitute a unique perturbation of both metal-ion and proton gradients that bacteria would attempt to counteract. Polyether ionophores can also affect eukaryotic cells, and while some compounds are used extensively as oral antiparasitic agents in the agricultural industry, the development of ionophore antibiotics for human use has not been pursued due to the risk of toxicity. Clearly, this is a multifaceted situation, however, the current best interpretation is that many organisms can tolerate polyether ionophores and that dosing schedules can be adapted to increase compliance.^13^ Despite the extensive use in animals, there is no evidence of cross-resistance between polyether ionophores and other antibiotics, which is interesting in relation to antimicrobial resistance (AMR).^14^

We hypothesize that exploitation of differences in membrane composition between mammalian and bacterial cells can be leveraged to prepare ionophores with acceptable therapeutic indices to be employed in certain clinical scenarios to treat resistant bacterial infections in humans. Recently, as an initial milestone towards this end, we reported on synthetic polyether ionophores with enhanced inhibitory selectivity for bacterial cells vs. mammalian cells compared to a series of benchmarked natural products.^15^ In the present study, we make a systematic comparison of nine members of the natural polyether ionophores (Fig. S1). While many of these agents have been subject to earlier investigations for their antibiotic properties, this data is often generated under different experimental conditions which we suspected might confound similarities (or differences) between the compounds. Furthermore, we demonstrate for the first time the antimicrobial properties of this class of compounds against bacterial biofilms and persister cells. Our data demonstrate surprising differences between members of this family of antibiotics and furthermore uncover inhibitory profiles for selected compounds that prompt investigations in preclinical infection models and for further SAR-investigations.

## Results and Discussion

### Polyether ionophores are effective across a panel of *Staphylococcus aureus* strains from bloodstream infections

To immediately establish a clinical relevance for investigating the ionophores as antibacterial compounds against *Staphylococcus aureus* (*S. aureus*), we determined the minimum inhibitory concentration (MIC) and the minimum bactericidal concentration (MBC) for several natural product ionophores against both methicillin-sensitive (MSSA) and methicillin-resistant (MRSA) *S. aureus* clinical isolates (Table 1). Two observations stand out from these experiments: 1) each ionophore are equally effective across all tested strains, and 2) all ionophores have a large difference between relatively low MIC values and MBC values that were not measurable in the tested range making the compounds bacteriostatic. A similar activity profile was observed with vancomycin used as the control. The only exception was calcimycin, a structurally atypical polyether ionophore.^16^

**Table 1.**
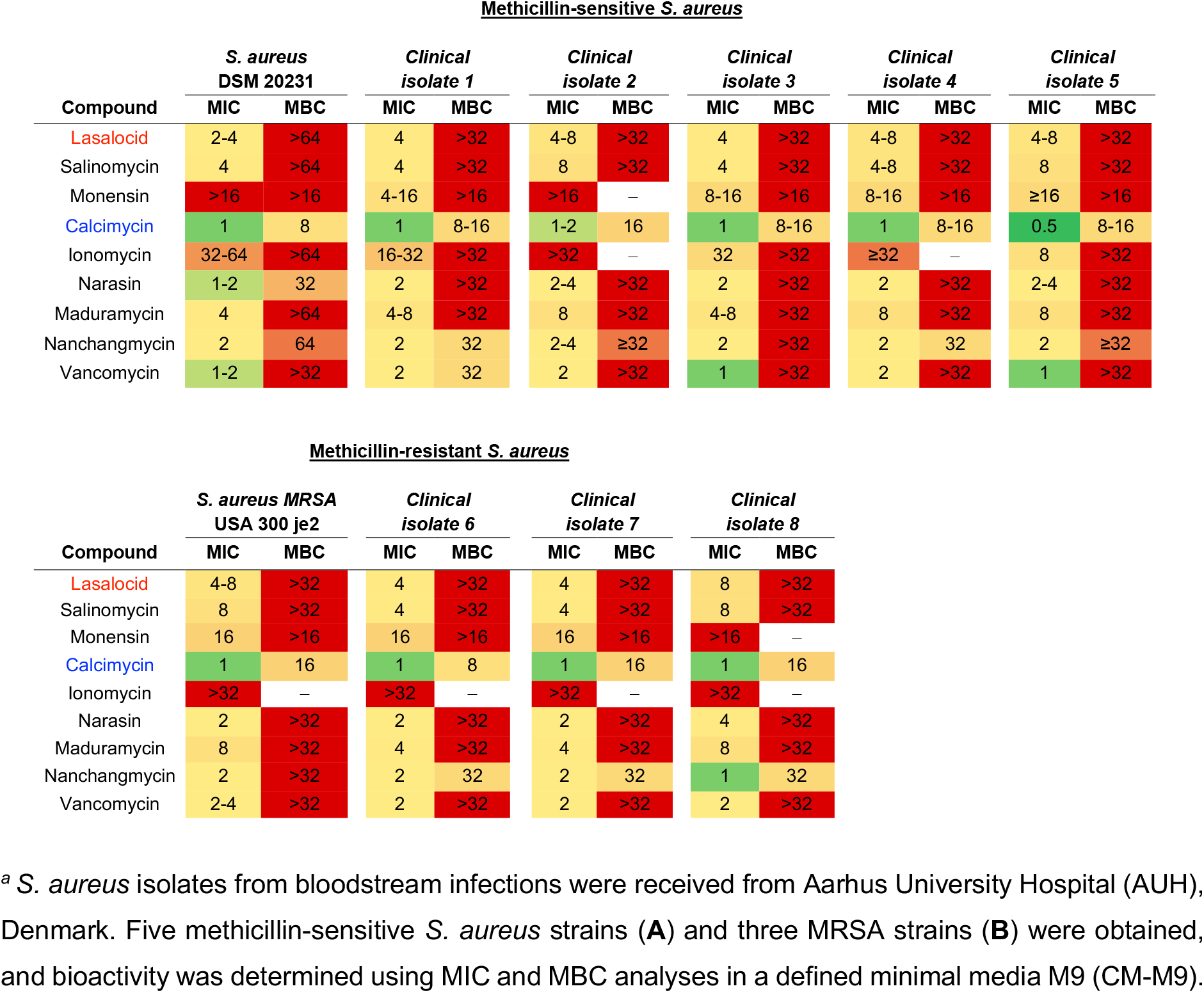
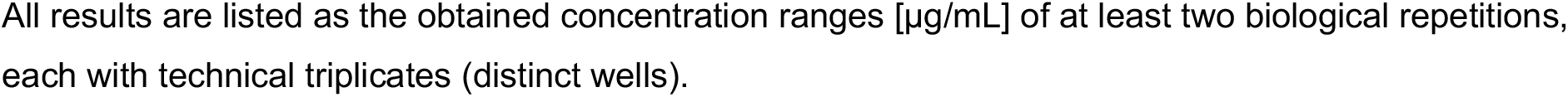
Polyether ionophore bioactivity in methicillin-sensitive *S. aureus* and methicillin-resistant *S. aureus* (MRSA) strains.^*a*^.

**Table 2.** MIC of polyether ionophores in *S. aureus* in the presence of different growth media.^*a*^.

| Compound | MIC [µg/mL] in *S. aureus* DSM 20231 |  |  |  |  | Maximum fold-change |
| --- | --- | --- | --- | --- | --- | --- |
|  | LB | MHB | CA-MHB | MHB + NaHCO<sub>3</sub> (25 mM) | CM-M9 |  |
| Lasalocid | 1-2 | 2 | 1 | 8-16 | 2-4 | 16 |
| Salinomycin | 0.5-1 | 2 | 1 | 4 | 4 | 8 |
| Monensin | 4-8 | 4-8 | 2 | 4 | >16 | ≥8 |
| Calcimycin | 0.06-0.25 | 0.13-0.25 | 1 | 1 | 1 | 16 |
| Ionomycin | 4-16 | >32 | >32 | ≥32 | ≥32 | ≥8 |
| Narasin | 0.25-0.5 | 0.5-1 | 0.25 | 1-2 | 1-2 | 8 |
| Maduramycin | 2 | 8-16 | 2 | 4 | 4 | 8 |
| Nanchangmycin | 2 | 4-8 | 4 | 1-2 | 2 | 4 |
| Vancomycin | 1 | 0.5-1 | 1-2 | 2-4 | 1-2 | 8 |

<sup>a</sup> The MIC determination was performed in technical triplicates and biological replicates in 96-well plates in accordance with CLSI recommendations.

We conducted these experiments in a defined minimal media CM-M9 (M9 salts with Ca^2+^ and Mg^2+^ adjusted to physiological level) and, serendipitously, noticed a large discrepancy between these values and those previously reported using type-strains in Lysogeny Broth (LB).^15^ Due to the rather broad mechanistic role of polyether antibiotics as ion transporters, we suspected that medium composition might play a significant role. Consequently, we determined the MIC values of our polyether ionophore panel in a small set of commonly used media: Lysogeny broth (LB), Mueller Hinton broth (MHB), cation-adjusted Mueller Hinton broth (CA-MHB), and a defined minimal media M9 (CM-M9). When compared to vancomycin – whose activity varied from 0.5-4 µg/mL dependent on medium composition (maximum 8-fold change) – some of the ionophores appeared more sensitive to change (more than 8-fold). Apart from maduramycin and nanchangmycin, the ionophores had the lowest MIC value in LB and the highest in CM-M9. A similar trend has been reported for calcimycin and ionomycin against *Bacillus subtilis*.^17^ We also determined the MIC in MHB supplemented with sodium bicarbonate (25 mM) to approach more clinically relevant screening conditions, because supplementation with sodium bicarbonate has been shown to improve *in vitro* to *in vivo* translatability of MHB.^6^ The MIC values mirrored those determined for the CM-M9 medium. Because the chemically defined CM-M9 allow for much better control over the individual cation concentration we chose to proceed with this in the subsequent experiments.

### The antibiotic properties of ionophores are affected by cation concentration

Based on the ionophore’s fluctuation in inhibitory potency observed between different media and their general role in ion transport, we hypothesized that cation concentrations would modulate their activity. This prompted us to systematically investigate the effect of changing the concentration of Na^+^, K^+^, Mg^2+^, Ca^2+^ in a full factorial design. Experimentally, the cation concentrations were set to vary within physiologically relevant ranges in a CM-M9 based medium. The ranges were set as follows: Na^+^ (65-125 mM), K^+^ (5-25 mM), Mg^2+^ (0.05-2 mM), and Ca^2+^ (0.1-2.5 mM) and we used growth as response upon challenge with ionophores at 0.25x, 1x, or 4xMIC (Fig. 1A). To validate the experimental setup, we also tested daptomycin, which should display calcium-dependence, and vancomycin which we did not expect to be influenced by changes in cation concentration within the set ranges. Indeed, daptomycin was potentiated with increasing calcium concentration and - to a smaller extent - increasing magnesium concentration. Vancomycin inhibition showed a slight inverse correlation with sodium concentration, but generally negligible cation dependency as expected (Fig. 1A). In comparison to daptomycin, the ionophore’s inhibitory effects were generally less impacted by changing cation concentrations and fell into two broad groups: calcimycin and ionomycin inhibition correlated with *increasing* calcium concentrations and, interestingly, *decreasing* magnesium concentrations, whereas lasalocid, salinomycin, narasin, and maduramycin inhibition correlated with increasing sodium and decreasing potassium concentrations. Calcimycin has also been reported to drastically lower the tolerated amount of added iron, manganese, and calcium to *B. subtilis*.^17^ The data largely reflect the affinity towards either monovalent (salinomycin, monensin, narasin) or divalent cations (calcimycin, ionomycin) that have been reported historically,^18^ although it is also clear that each compound has a unique profile.

**Fig. 1.**
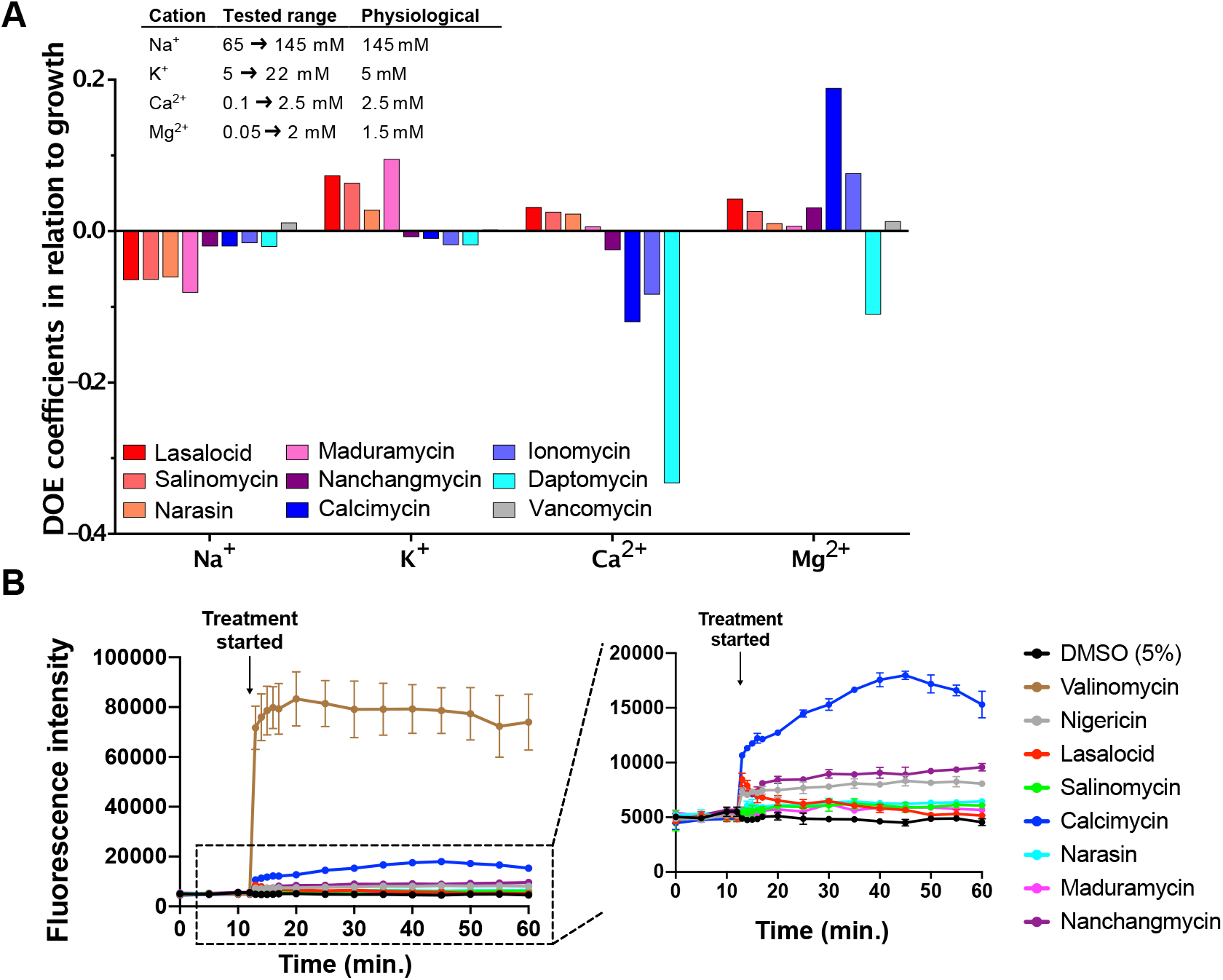
Cation-dependency of polyether ionophores. **(A)** A full factorial design of experiment (DOE)- approach was used to study how ionophore activity depends on cation-concentrations (Na^+^, K^+^, Ca^2+^ and Mg^2+^). The concentration of the four cations was chosen as factors, and the percentage change in growth compared to a growth control as response. *S. aureus* was treated with the compounds at 0.25xMIC concentration (1xMIC concentration for daptomycin) in presence of cations in the concentration ranges listed in the figure. Bacterial growth was evaluated after overnight treatment, and MODDE Pro was used for data processing. Daptomycin and vancomycin were included as positive and negative controls, respectively. Results are average of technical triplicates. **(B)** The ionophores effects on membrane potential were studied using the membrane-potential sensitive dye 3,3-dipropylthiacarbocyanine iodide (DiSC_3_(5)). *S. aureus* was loaded with 1 µM DiSC_3_(5) (1%) and after the dye was stabilized, the culture was treated with 10xMIC concentration (CM-M9 media). Valinomycin [20 µg/mL] and nigericin [10 µg/mL] were included as positive and negative controls, respectively. Results are average of technical triplicates (distinct wells), and bars are mean ±s.d. (n=3).

As a consequence of their dual ion-transport and membrane binding activities, polyether ionophores may acutely disrupt cellular membrane potential. This effect could be the cause of both anti-bacterial and cytotoxic effects. To probe the effect of ionophores on membrane potential, we used the dye 3,3-dipropylthiacarbocyanine iodide (DiSC_3_(5)). DiSC_3_(5) aggregates within the bacterial membrane causing self-quenching of its fluorescence but upon disruption of the membrane electrical potential (ΔΨ), the dye is released into the medium accompanied by an increase in fluorescence intensity. We tested eight naturally occurring polyether ionophores at 10xMIC in this setup (Fig. 2B). As a positive control, we used the pore-forming K^+^ ionophore valinomycin, which is known to dissipate the ΔΨ. In comparison to valinomycin, the polyether ionophores did not dissipate ΔΨ as the fluorescence intensity remained stable for 60 min (Fig. 2B). Only calcimycin caused a slight increase in fluorescence intensity, although much less than valinomycin. Work by Farha et al. have shown that DiSC_3_(5) can indirectly probe the proton gradient (ΔpH) across the membrane because bacteria will respond to a decreasing ΔpH by increasing ΔΨ, thereby leading to decreased fluorescence readout by DiSc_3_(5).^19^ This latter effect was reported previously for nigericin, but we did not observe any significant response for the ionophores tested here. In conclusion, our data indicate that the ionophores do not disrupt the membrane potential in any significant way. Another study by Farha et al. showed that the drastic changes in potency of antibiotics at increased sodium bicarbonate levels are caused by a selective dissipation of the ΔpH.^7^ They also showed that dissipators of ΔΨ (valinomycin, I1, I2, I3) act synergistic with bicarbonate. If the ionophores did in fact function as potent dissipators of the membrane potential, we should see a more pronounced effect on the measured MIC values when adding bicarbonate.

**Fig. 2.**
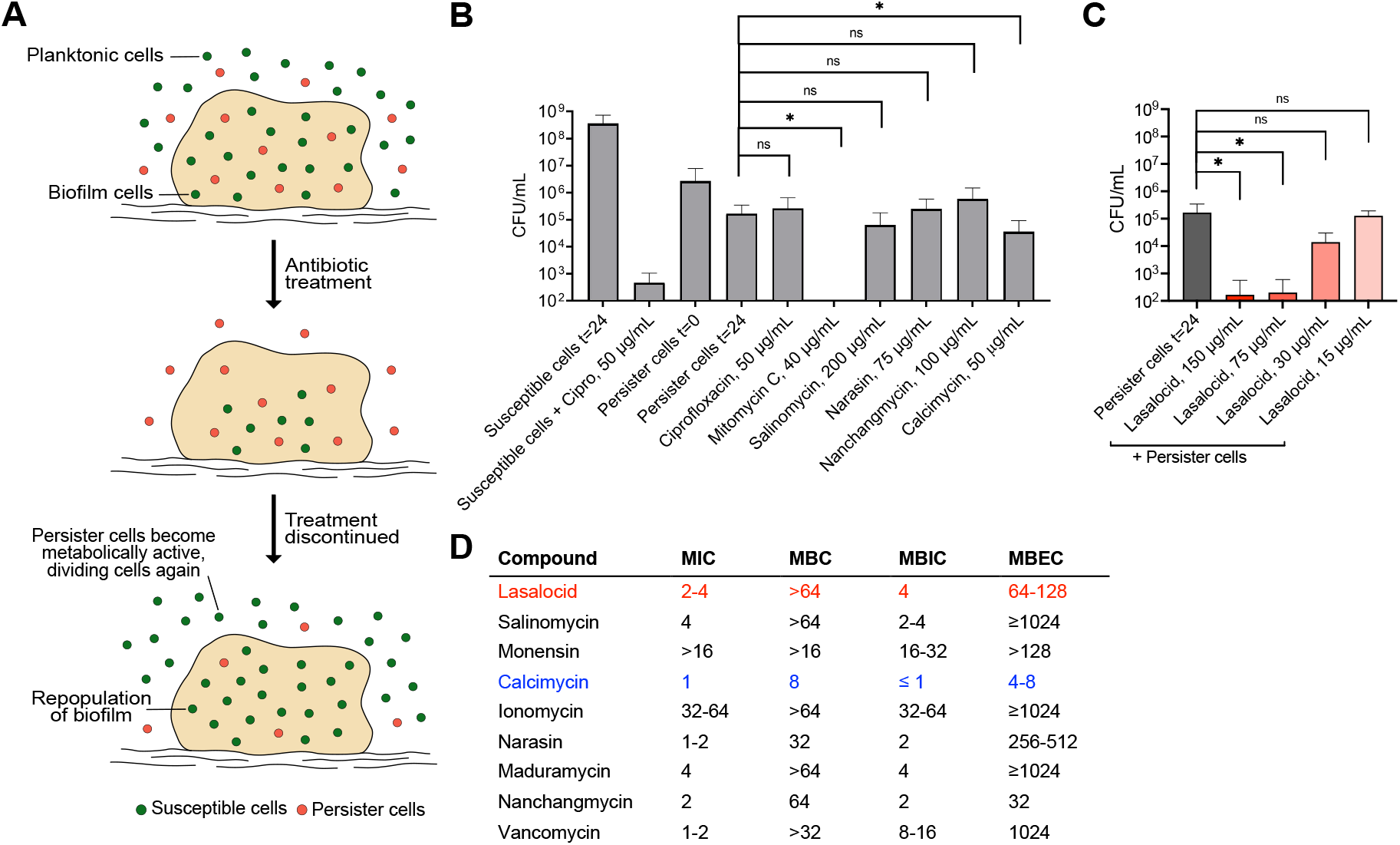
Polyether ionophore action on *S. aureus* persister cells and biofilms. **(A)** Illustration showing the lack of antibiotic effect against persister and biofilm cells. **(B)** Persister cell killing with 50xMIC concentration (the respective concentrations are listed in the figure). Persister cells were produced by transferring growing, susceptible bacteria into a modified CM-M9 medium without available carbon sources. Persister cells were reverted to susceptible cells when transferred into a nutrient rich media (susceptible cells t=24). Ciprofloxacin (50 µg/mL) and mitomycin C (40 µg/mL) were included as negative and positive controls, respectively. Bars are mean ± s.d. (n=9). **(C)** Persister cell killing with lasalocid at 5x, 10x, 25x and 50xMIC concentrations. Bars are mean ± s.d. (n=9). **(D)** Biofilm inhibition and eradication. A *S. aureus* biofilm was grown for 24 hours before antibiotic treatment. Minimum biofilm inhibition concentration (MBIC) was determined after 24 hours of treatment, and minimum biofilm eradication concentration (MBEC) was determined after 72 hours of biofilm recovery. Lasalocid (red) and calcimycin (blue) are highlighted for clarity. All results shown are averages of biological triplicates, each with technical triplicates.

**Fig 3.**
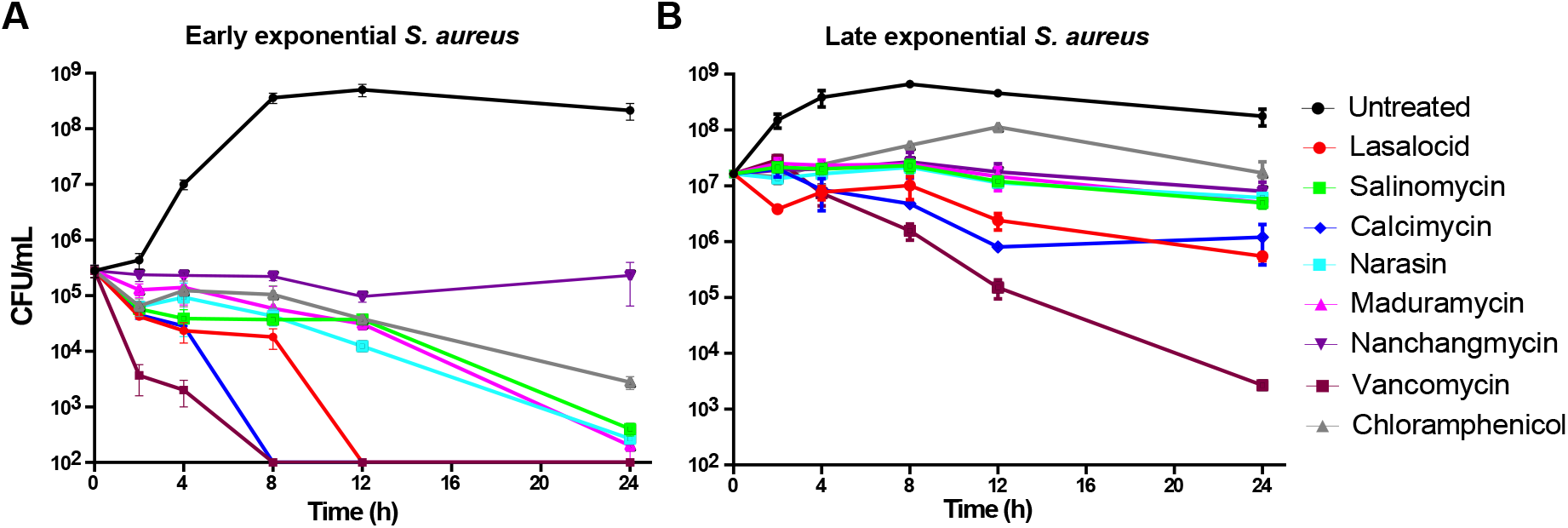
Time-dependent killing of *S. aureus. S. aureus* was grown to early (**A**) and late (**B**) exponential phase and treated with ionophores at 10xMIC (in CM-M9 media) concentration for 0, 2, 4, 8, 12 and 24 hours where cells were harvested, and CFU/mL was enumerated (cut-off 10^2^ CFU/mL). Vancomycin (10xMIC of 2 µg/mL) and chloramphenicol (1xMIC of 16 µg/mL) were included as bactericidal and bacteriostatic control, respectively. Individual data points are mean±s.d. (n=3) of technical triplicates (distinct wells). The figure is a representative of two biological replicates.

### Selected polyether ionophores kill bacterial persisters and biofilms

We next investigated antibacterial effects beyond simple early exponential growth. Two of the most substantial clinical challenges in relation to infections are the formation of biofilms and persister cells, both of which are primary causes of chronic and recurrent infections.^20–23^ Upon antibiotic treatment, bacterial cells can transform into metabolically inactive persister cells, making them highly resilient to treatment and in this way ‘escape’ the antibiotic (Fig. 2A). When antibiotic treatment is discontinued, the persister cells eventually transition back to metabolically active cells creating recurrent infections. Biofilms, which typically grow on surfaces of medical implants or host tissue, are characterized by an increased tolerance to antibiotic treatment. The reason behind this increased tolerance is attributed to lower growth rate, decreased diffusion of antibiotics, higher tolerance to oxidative stress, and a larger subpopulation of persister cells (Fig. 2A).^24^ To investigate the effects of polyether ionophores on metabolically inactive persister cells, we transferred *S. aureus* cells to CM-M9 medium with the carbon source omitted - thereby preventing any growth and mimicking the subpopulation left after antibiotic treatment – and incubated with antibiotics for 24 hours before determining the viable cell count. We used ciprofloxacin (50 µg/mL) as negative control (no inhibitory activity against persister cells) and the DNA-crosslinker mitomycin C (40 µg/mL) as the positive control for killing persister cells. When tested at 50xMIC the ionophores displayed very different inhibitory potential in this setting (Fig. 2B). While most compounds were completely inactive, calcimycin showed a modest effect on the persister cells at 50xMIC, but lasalocid showed a concentration-dependent inhibitory effect on the persister cells and a remarkable potency at >75 µg/mL. Next, using a 96-well assay based on peg-lids, the minimum biofilm inhibition concentration (MBIC) and minimum biofilm eradication concentration (MBEC) were determined for the ionophores (Fig. 2D). For most of the compounds, the MBIC mirrored the MIC, while the MBEC was much higher than the MBC, which is in agreement with previous reports on *S. epidermidis* and *S. aureus* on some of the ionophores.^25,26^ However, lasalocid, nanchangmycin, and calcimycin stood out with MBEC values similar to their MBC, indicating that under these test conditions the biofilm matrix and the persister phenotype in these biofilms do not sufficiently protect the bacteria against the antimicrobial action of the ionophores, but the proximate cause for this is still unclear.

### Time-dependent killing of *S. aureus*

After establishing the dependencies of medium composition and cation concentration, we challenged *S. aureus* with the ionophores (10xMIC) in a time-dependent fashion and compared ionophore activity to chloramphenicol and vancomycin. In an early exponential culture (5 × 10^5^ CFU/mL), all ionophores displayed bactericidal behavior except for nanchangmycin. Lasalocid and calcimycin caused a rapid decline in colony forming units (CFU) that dropped below our level of detection within just 12 hours. The same two compounds also showed the largest effect in a late exponential culture (2 × 10^7^ CFU/mL). This also reflects the rather large difference in MIC and MBC that we observe for many of the ionophores. In conclusion, when the effects in the biofilm and persister assays and time-dependent killing, are viewed in combination, nanchangmycin, lasalocid, and calcimycin stand out as promising for further investigation as antibiotic leads.

### Genes affecting ionophore susceptibility

When assessing the potential of polyether ionophores as antibiotic leads an important aspect to address is the frequency of resistance, especially considering that many ionophores are used in agriculture. In addition, this can also provide insights into an antibiotic’s mechanism of action as mutations conferring resistance can reveal the compound’s target. The producing organisms of the ionophores often contain genes conferring self-resistance. In the case of the ionophore tetronasin, Linton et al. showed that two genes encoding a ATP-dependent efflux system in *S. longisporoflavus* conferred resistance to tetronasin when transferred to otherwise susceptible *S. lividans* and *S. albus*.^27^ Similarly, the biosynthetic gene clusters associated with monensin (BGC0000100), lasalocid (BGC0000087), and nanchangmycin (BGC0000105) found in the MIBiG database^28^ all contain predicted efflux systems indicating that this is the dominant resistance mechanism among producing strains.

To investigate this in a human pathogen, we examined resistance against selected ionophores, first – unsuccessfully – in the standard single-step challenge of a dense inoculum on agar plates containing 1xMBC, and next by a sequential passaging method where an *S. aureus* culture was treated with sub-inhibitory concentrations every day for 4 weeks (Fig. 4). In this type of experiment, as the bacteria become less susceptible with time, and perhaps resistant, the MIC value will increase. However, after 25 days of continuous treatment, the *S. aureus* culture was barely affected by the exposure. For lasalocid, salinomycin and calcimycin, only a 4-fold increase in the tolerated concentration was observed which is comparable to vancomycin that we used as control. In comparison, resistance to ciprofloxacin developed rapidly as would be expected. These data mirror prior observations in the literature that resistance towards polyether ionophores is not easily achieved and that their frequency of resistance is very low considering their extensive use.^29^ Interestingly, the *S. aureus* culture treated with nanchangmycin was slightly more affected after the first 25 days of treatment with a 16-fold change in inhibitory concentration and this treatment was therefore continued. After 40 days of continuous treatment, susceptibility was decreased resulting in an apparent 64- to 128-fold increase in MIC. When isolating single colonies to determine the final MIC value of the resistant strains, a modest increase of only 4- to 16-fold increase in MIC compared to the wildtype *S. aureus* strain was however observed (Fig. 4B). The nanchangmycin-resistant strains were checked for cross-resistance towards the other three ionophores as this could indicate whether a shared resistance mechanism was involved. However, none of the isolates showed any difference in tolerance to lasalocid, salinomycin or calcimycin thereby ruling out a common acquired resistant phenotype. We sequenced the genomes of three resistant strains as well as the wild type *S. aureus* strain, which revealed several genes with mutations common among all three mutants and absent in the wildtype (see Table S2). Among these were the gene encoding TrkH (WP_000021864.1) involved in potassium uptake; the gene encoding MspA (WP_001161085.1), a protein hypothesized to be membrane stabilizing;^30^ and the gene encoding the transcriptional regulator SarV (WP_000066900.1) that is involved in the autolysis of *S. aureus*.^31^ Common for strain 1 and 3 were a large number of mutations in both *mspA* and *sarV*, while strain 2 only showed one mutation in each. These results matches previous findings that acquired ionophore-resistance is linked to efflux or decreased permeability,^32,33^ but the proximate cause remains elusive. To gain additional insight into the mechanism of action of the ionophores, we screened the Nebraska Transposon Mutant Library (NTML), a mutant library in an MRSA strain with transposons inserted into 1,920 individual non-essential genes, for increased sensitivity against lasalocid, salinomycin, calcimycin, and nanchangmycin in an agar plate assay. Here, we primarily found hits with perturbations in the electron transport chain to be more sensitive to the ionophores (see Table S2). Measuring the MICs of the mutants in a liquid-based assay revealed that inactivation of *aroC* (encoding chorismate synthase) or *hemB* (encoding delta-aminolevulinic acid dehydratase) had the biggest effect on ionophore sensitivity, while mutations inside genes of the quinol oxidase subunits I, II or III (*qoxABC*), NADH:ubiquinone reductase (*ndh2*) or protoheme IX farnesyltransferase (*cyoE*) gave 2-fold decreases in the MIC values of lasalocid, salinomycin, and nanchangmycin. None of the mutations influenced calcimycin sensitivity in a liquid setting. We have previously observed that many of these transposon mutants have a reduced membrane potential^34^ and *hemB* has also been associated with small colony variants.^35^

**Fig. 4.**
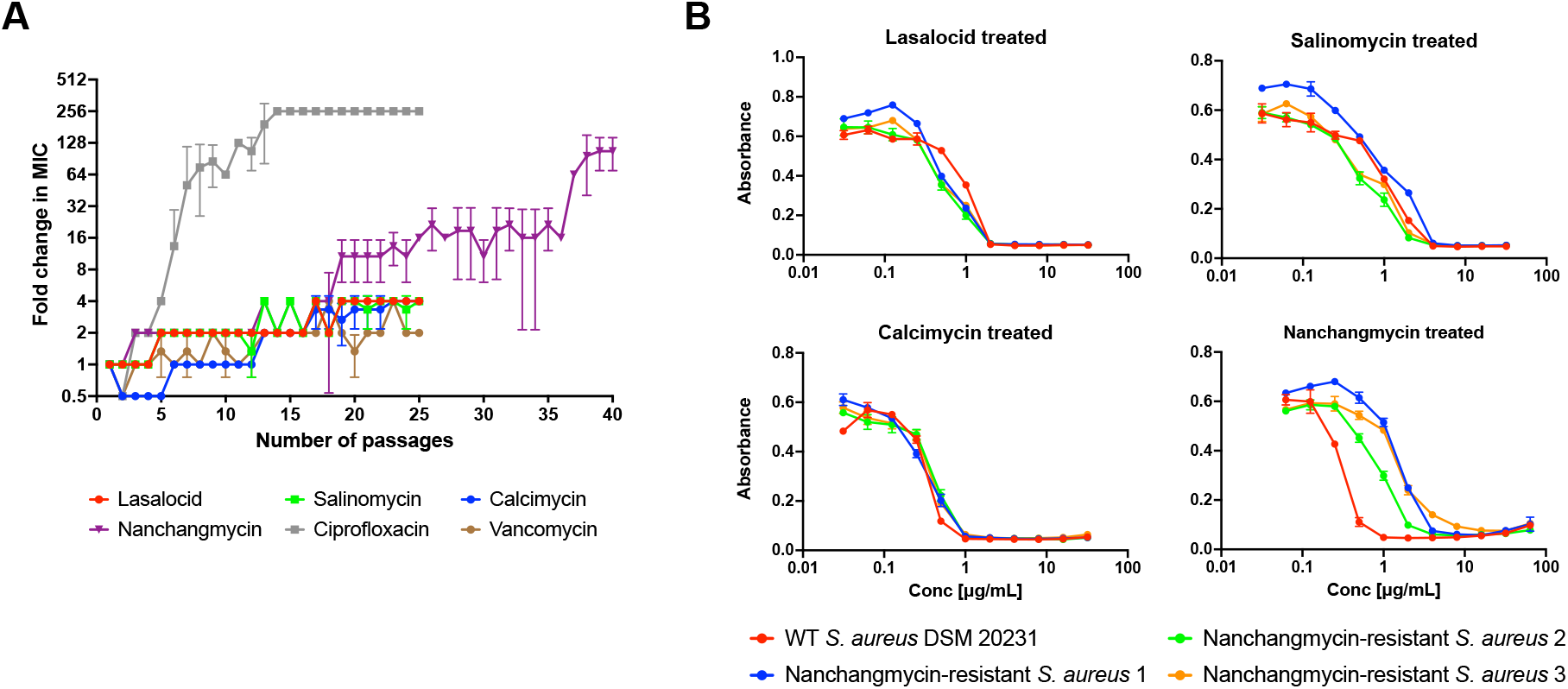
Resistance studies. **(A)** Resistance towards ionophores was attempted developed with a sequential passaging method. *S. aureus* was treated with five ionophore concentrations (0.25xMIC to 4xMIC), and MIC was determined every day. The culture growing in presence of highest concentration (0.5x new MIC) was passed on to following day and treated with concentrations relative to the new MIC value. This was continued for 25-40 days. Ciprofloxacin and vancomycin were included as positive and negative controls, respectively. Data points are mean ± s.d. (n=3). **(B)** The three nanchangmycin-treated cultures on day 40 in (A) were subjected to MIC analysis to confirm resistance and checked for cross-resistance towards the other three ionophores. Wildtype *S. aureus* DSM 20231 was included as control strain. Data points are mean ± s.d. (n=3), and results are a representative of biological duplicates.

### Polyether ionophores synergize with polymyxin B to target Gram-negative pathogens

Polyether ionophores are generally regarded as inactive against Gram-negative pathogens. Although their abilities, that we have documented in this study, to target drug-resistant, Gram-positive clinical isolates, as well as persister populations and bacterial biofilm, is undoubtedly appealing, the lack of Gram-negative activity is a clear limitation of this compound class. It is likely that polyether ionophores cannot efficiently penetrate the outer membrane of Gram-negative bacteria and that this is the cause of their low activity. Although the uptake efficiency, to the best of our knowledge, has not been experimentally determined for all of the polyether ionophores in Gram-negative strains, Guyot et al. showed that calcimycin bound to *Escherichia coli* but never entered the cells.^36^ The lack of penetration is also supported by studies employing *E. coli* mutant strains having increased outer membrane permeability, that remain sensitive to polyether ionophores or studies showing that the polymyxin derivative PMBN sensitizes *E. coli* to calcimycin.^15,37^ We hypothesized that combinations with compounds such as polymyxin B, which specifically target the outer membrane and is known to increase permeability of other antibiotics, might allow all of the ionophores to act against Gram-negative pathogens including *E. coli* and we therefore performed a systematic study of these effects. None of the ionophores had any stand-alone effect against *E. coli* (or *Pseudomonas aeruginosa* and *Acinetobacter baumannii*, Fig. S2), but salinomycin (FIC 0.25), calcimycin (FIC 0.125) and nanchangmycin (FIC 0.25) all lowered the MIC value for polymyxin B in a dose-dependent manner (Fig. 5). Conducting the same experiment with an antibiotic with an intracellular target (kanamycin, ribosome) indicates that the outer membrane is in fact the only thing preventing the ionophores from exerting their effect on Gram-negative bacteria. Interestingly, lasalocid did not afford clear synergies with polymyxin B, an observation that we are currently not capable of explaining. Collectively, these data suggest that inclusion of polyether ionophores in antibiotic cocktails should be systematically tested in pre-clinical settings.

**Fig. 5.**
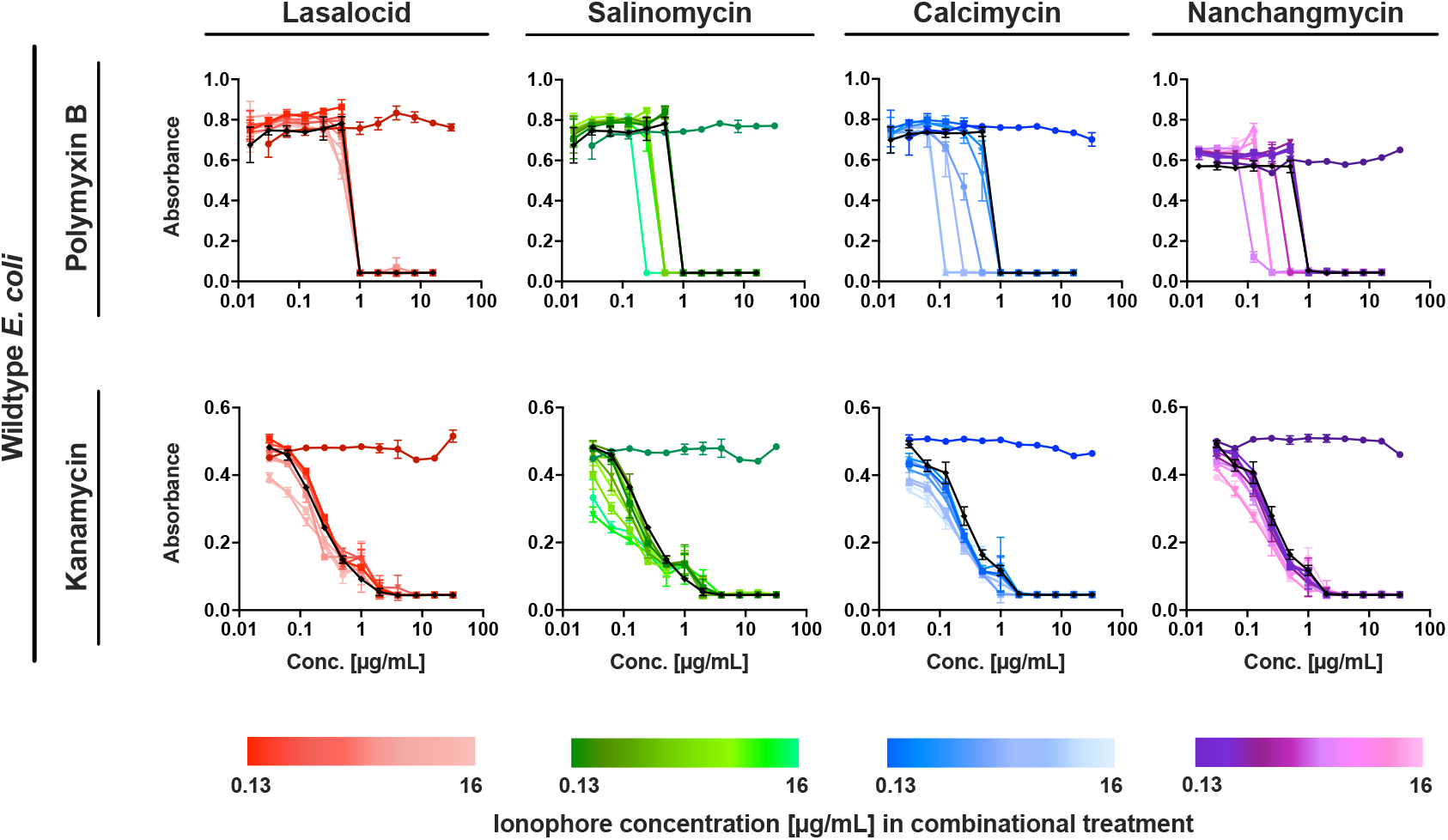
Combinatorial treatment in wildtype *E. coli. E. coli* were treated with either polymyxin B or kanamycin (black curves) and in combination with selected ionophores which were added as a fixed dosage (concentration range 0.13-16 µg/mL). The antibiotics were tested as 2-fold serial diluted up to 16 µg/mL (polymyxin B) or up to 32 µg/mL (kanamycin). The ionophores were tested as single treatments (lasalocid: dark red; salinomycin: dark green; calcimycin: dark blue; nanchangmycin: dark purple) as negative controls. The results are representative of two biological replicates, and data points are mean ±s.d. (n=3).

## Conclusion

In the work reported here, we have demonstrated that polyether ionophores, despite their joint, canonical characteristics as ion-transporters, are in fact quite heterogeneous with respect to their antibacterial properties. The compounds’ dependencies on cation concentrations are distinct, and some members display the ability to eradicate bacterial biofilms and inhibit persister populations. We show that resistance to polyether ionophores is difficult to achieve and does not result in cross-resistance within the ionophore family. Very recently, Zhu et al. reported equally promising persister and biofilm potency for the ionophore nigericin,^38^ a compound we had originally omitted due to its low solubility in our carrier DMSO. They demonstrated *in vivo* efficacy against S. aureus in a mouse model, emphasizing the potential of the ionophores as antibiotics.

Finally, we show that combinations with polymyxin B can expand the antibiotic activity of the ionophores into also covering Gram-negative bacteria. This is particularly interesting given the increasing challenges associated with drug-resistant, Gram-negative pathogens.

## Acknowledgements

This project has received funding from the European Research Council (ERC) under the European Union’s Horizon 2020 research and innovation programme (grant agreement number 865738). Financial support from Independent Research Fund Denmark (grant number 9040-00117B) is acknowledged.

## Materials and methods

### Bacterial strains and culture conditions

*Staphyloccocus aureus* DSM 20231 was studied in most of the conducted experiments. *Escherichia coli* BW25113 (obtained from the Keio Collection, Coli Genetic Stock Center, Yale University), *Pseudonomas aeruginosa* DSM 19880, and *Acinetobacter baumannii* DSM 300007 were included in combinatory treatment experiments. *S. aureus* isolates from bloodstream infections were obtained from the Department of Clinical Microbiology, Aarhus University Hospital.

All strains were cultured in a defined minimal M9 (CM-M9) media unless otherwise stated. This media contains M9 broth (purchased at Sigma-Aldrich with product number 63011, containing NH_4_Cl (18.7 mM), Na_2_HPO_4_ (42.3 mM), KH_2_PO_4_ (22 mM) and NaCl (8.6 mM)), glucose (1%), casamino acids (1%), CaCl_2_.2H_2_O (0.1 mM), MgSO_4_.7H_2_O (2 mM), thiamine hydrochloride (1 mM), nicotinamide (0.05 mM), and trace metal solution TMS3 (1 mL/L), pH 7.4.

### Polyether ionophores

The chemical structures and selected properties of the polyether ionophores used in the current study can be found in Fig. S1. The compounds was obtained from the following sources: Calcimycin (Alomone labs, A-600), Ionomycin (Alomone labs, I-700), Narasin (Sigma, N1271), Salinomycin (Toku-E, S002), Monensin (Toku-E, M083), Maduramycin (Sigma, 34069), Nanchangmycin (Sigma, SML2251), Lasalocid was prepared by isolation of its sodium salt from commercially available veterinary premix AVATEC, followed by acidic extraction with H_2_SO_4_ and crystallization from ethanol but can also be obtained from commercial suppliers (Sigma, 73996).

### Minimum inhibitory concentration (MIC) analysis

MIC analysis was performed by a broth microdilution method in 96-well microtiter plates in accordance with CLSI recommendations. Briefly, a culture was grown aerated overnight (16-18h, 140 rpm, 37°C) and adjusted to a final density of 5×10^5^ CFU/mL. Compound stocks were prepared in 2-fold serial dilutions in DMSO (final conc. <5%). Bacteria were treated with compounds overnight (20-24h, 37°C), and growth was measured as absorbance at 600 nm using a Tecan Nanoquant infinite M200Pro plate reader. The absorbance value correlating to a difference seen by the unaided eye was determined to absorbance 0.07, and measurements above this value were deemed as growth when determining the MIC value.

### Minimum bactericidal concentration (MBC) analysis

MBC analysis was performed after the MIC analysis of all concentrations without detectable growth. Cells were pelleted in the 96-well microtiter plates (15 min, 3000 rpm), and pellets were resuspended in 0.9% NaCl. 10 µL of the suspension was spotted onto LB agar and allowed to dissolve into the agar. A concentration with visible growth was included as positive control. Agar plates were incubated overnight (16-24h, 37°C) and checked for growth. A bactericidal effect is represented by a 3 log10 reduction in density corresponding to a 99.9% reduction. Thus, spots with less than 5 colonies indicate a bactericidal effect and are stated as the MBC value.

### Biofilm assay

A culture was grown aerated overnight (16-18h, 37°C, 140 rpm) and adjusted to 1×10^7^ CFU/mL. The adjusted culture was inoculated with a peg-lid (30 min, 37°C) to adhere biofilm cells onto pegs. The peg-lid was then transferred to fresh media and allowed to grow overnight (24h, 37°C) before treatment. Biofilm cells were treated for 24 hours (37°C) in CM-M9 with compounds (dissolved in DMSO) in a 2-fold serial dilution prepared in medium to lower DMSO concentration. The peg-lid was washed twice in 0.9% NaCl and sonicated for 10 min in fresh medium, and biofilm cells were allowed to recover (72h, 37°C). The minimum biofilm inhibitory concentration (MBIC) was determined after 24 hours of treatment, and the minimum biofilm eradication concentration (MBEC) was determined after 72 hours of recovery, both with absorbance measurement at 600 nm and visual inspection.

### Persister cell assay

A culture was grown aerated overnight (22-25h, 37°C, 140 rpm) in tryptic soy broth (TSB), diluted 1:1000 in fresh TSB and incubated overnight a second time. The culture was adjusted to OD_600_ 1.0 and washed using a modified CM-M9 medium (without glucose and casamino acids). In a 96-well microtiter plate, compounds were dispensed in modified CM-M9 at 50xMIC concentration, and the adjusted culture was added yielding a final density of OD_600_ 0.1. The plate was incubated overnight (20-24h, 37°C), and pellets were washed by pelleting (10 min, 14,000 rpm) in a tabletop centrifuge in eppendorf tubes to remove compounds. The cells were finally 10-fold serial diluted in modified CM-M9, and 10 µL from dilution 10^0^ to 10^7^ was spotted onto LB agar. On the following day, colonies were counted for CFU/mL enumeration.

### Time-dependent killing

For early-exponential phase, a culture was grown aerated overnight (16-18h, 37°C, 140 rpm) in CM-M9 media, adjusted to 1×10^5^ CFU/mL and grown to early exponential phase (3h, 37°C, 140 rpm) before treatment. For late-exponential phase, a culture was grown aerated overnight (16-18h, 37°C, 140 rpm) in CM-M9 media and used directly for treatment. In both cases, the cultures were diluted 1:10 with CM-M9 and treated with compounds in a 96-well format at 10xMIC concentration (CM-M9 media). At six chosen timepoints (0, 2, 4, 8, 12 and 24 hours), samples were collected for CFU/mL enumeration. Here, the cells were washed (15 min, 3000 rpm) in 0.9% NaCl in the 96-well plates, 10-fold serial diluted in 0.9% NaCl, and 10 µL from each dilution (10^0^ to 10^6^) was spotted onto LB agar. After incubation (16-24h, 37°C), CFU/mL was enumerated.

### Resistance development by single-step method

Agar plates were prepared with 1xMBC concentration of the desired compound by addition of compound (dissolved in DMSO) to 50-60°C LB agar. Meanwhile, a culture was grown aerated overnight (16-18h, 37°C, 140 rpm) and concentrated 20X by centrifugation (20 min, 4000 rpm), yielding a final density of ∼10^9^ CFU/mL. 100 µL of the dense culture was plated onto the agar plates containing compound, and plates were incubated for 24-72 hours (37°C) and checked for growth every day.

### Resistance development by sequential passaging

An overnight culture (16-18h, 37°C, 140 rpm) with density 1×10^8^ CFU/mL was diluted 1:40 and treated with five different compound concentrations (0.25x, 0.5x, 1x, 2x and 4xMIC) in a 96-well microtiter plate (one plate per technical triplicate). The plates were incubated overnight (22-24h, 37°C) and checked for growth. The new MIC value was determined with visual inspection, and 5 µL from the well with highest concentration with detectable growth (0.5x new MIC value) was passed on and treated with the five compound concentrations (0.25x to 4xMIC) relative to the new MIC value, diluting the culture 1:40. This was repeated for 25 days or until resistance was developed. Every day, an untreated growth control and medium blank were included.

### Isolation of genomic DNA and DNA sequencing

The genomic DNA (gDNA) was isolated from the nanchangmycin-resistant strains to perform DNA sequencing. For isolation of gDNA, the Monarch^®^ Genomic DNA purification kit was used (#T3010S) together with bead-beating with zirconia/silica beads. The procedure for the Monarch^®^ kit was followed with a few adjudgments, as followed. Overnight cultures (16-18h, 37°C, 140 rpm) of each resistant-strain and wt *S. aureus* DSM 20231 were grown to densities of ∼1×10^8^ CFU/mL and pelleted (1 min, 14,000 rpm). Pellets were resuspended in Tris-buffer and transferred to bead columns. Samples were beat beaten (40 sec, 6.0 m/s), beads were pelleted (10 min, 12,300 x g), and supernatant was stored in freezer overnight. The samples were then treated with enzymes: first with Lysozyme (20 mg/mL, 1h, 37°C), then with Proteinase K (25 µL, 2h, 56°C, 1400 rpm) and finally with RNase A (3 µL, 10 min, 56°C, 1400 rpm). The lysed samples were then processed as described in the Monarch^®^ protocol for gDNA binding, washing and elution. The samples were prepared for sequencing using the Nextera XT DNA Library Preparation Kit (Illumina) and paired-end sequenced (2 × 300 bp) on a MiSeq sequencer using the MiSeq Reagent kit V3 (Illumina). Geneious was used to trim the sequencing data (using BBduk) and then mapped onto the reported genome of S. aureus DSM 20231 (NZ_CP011526). All data has been deposited with GenBank under Bioproject number PRJNA877072 and accession numbers: CP104478 (untreated reference), CP104477 (resistant mutant 1), CP104476 (resistant mutant 2), CP104475 (resistant mutant 3).

### Design of experiment (DOE) approach

Using the DOE-approach, the ionophore’s bioactivity was studied in presence of different cation concentrations. To test the desired concentrations, a modified version of the CM-M9 medium was prepared containing NH_4_Cl (1 g/L), Na_2_HPO_4_ (7.5 g/L) and NaH_2_PO_4_ (3 g/L) instead of the M9 broth. The desired concentrations of NaCl, KCl, CaCl_2_ and MgSO_4_ could then be added. 17 different cation combinations wanted to be tested and thus, 17 different media were prepared.

A culture was grown aerated overnight (16-18h, 37°, 140 rpm) and diluted to a final density of 5×10^5^ CFU/mL. The bacteria were treated with three different compound concentrations (0.25x, 1x and 4xMIC) in the 17 different types of media in 96-well microtiter plates. Plates were incubated overnight (20-24h, 37°C), and growth was measured with absorbance measurements at 600 nm. A plate with media blanks was also prepared and these absorbance values were subtracted from the growth values giving the final growth responses. These values were plotted into the MODDE pro program which performed statistically analysis showing how each cation affects growth among other features.

### Fluorescence with 3,3’-dipropylthiacarbocyanine iodide (DiSC_3_(5))

The effect on membrane potential was studied with the fluorogenic probe 3,3’-dipropylthiacarbocyanine iodide (DiSC_3_(5)).

A culture was grown aerated overnight (16-18h, 37°C, 140 rpm), diluted to OD_600_ 0.1 and incubated to early-exponential phase (3h, 37°C, 140 rpm). The culture was washed (4000 rpm, 15 min, 4°C) three times in a sucrose buffer containing K_2_HPO_4_ .3H_2_O (10 mM), MgSO_4_ .7H_2_O (5 mM) and sucrose (250 mM), pH 7.0. After the final wash, the pellet was resuspended in the buffer supplemented with 0.1 M KCl to a final density of OD_600_ 0.3.

A culture plate was prepared in a black 96-well plate (Greiner bio-one 96-well µclear, black with clear bottom) where the diluted culture was loaded with 1% of fluorescent dye DiSC_3_(5), yielding a final concentration of 1 µM DiSC_3_(5). The dye was allowed to stabilize for 15 min in the dark before fluorescence was measured (excitation 620±10 nm; emission 685±10 nm; gain 2600, focal height 7.3 mm, top optic, and linear shaking of 500 rpm, 10s). In another black 96-well plate, compounds were dispensed as a 20X solution. Compounds were tested at 10xMIC concentrations, so stock solutions of 200xMIC were prepared in DMSO (final conc. 5%). After the dye was stabilized inside the bacterial membrane, the culture loaded with dye was transferred to the compound plate yielding the final concentration of 10xMIC. Fluorescence was measured for a total of 60 min. Controls of DMSO (5%) and buffer were included.

### Combinatorial treatments

For combinatorial treatments, the experiments were performed as stated in MIC analysis but with two compounds instead of one. The antibiotics were tested in a 2-fold serial dilution up to 16 µg/mL for polymyxin B and cloxacillin, or up to 32 µg/mL for kanamycin, and the ionophores were added as a fixed dosage for each serial dilution. 8 different combinations were made, testing the ionophores in the concentration range 0.125-16 µg/mL. The ionophores and antibiotics were also tested as single treatments as negative and positive controls, respectively.

### Screening of Nebraska transposon mutant library

The Nebraska Transposon Mutant Library is stored in glycerol in 96-well microtiter plates at - 80 °C. Material from the frozen stock was transferred with a Deutz 96 cryo-replicator from the 96-well microtiter plates onto Tryptic Soy Broth Agar plates supplemented with either 0.5 µg/mL lasalocid or salinomycin, 0.0625 µg/mL calcimycin or 0.125 µg/mL calcimycin, corresponding to 1/16 MIC of the MRSA JE2 wildtype strain in Tryptic Soy Broth (TSB). The plates were incubated at 37 °C for 24 h and visually inspected for lack of growth of individual mutants. MICs of individual mutants were performed in TSB in a 96-well format and determined by lowest concentration with no visible growth.

