## Supplementary material for "Polyether ionophore antibiotics target drug-resistant clinical isolates, persister cells, and biofilms": Sup. Fig S1-2 Sup. table S1-3

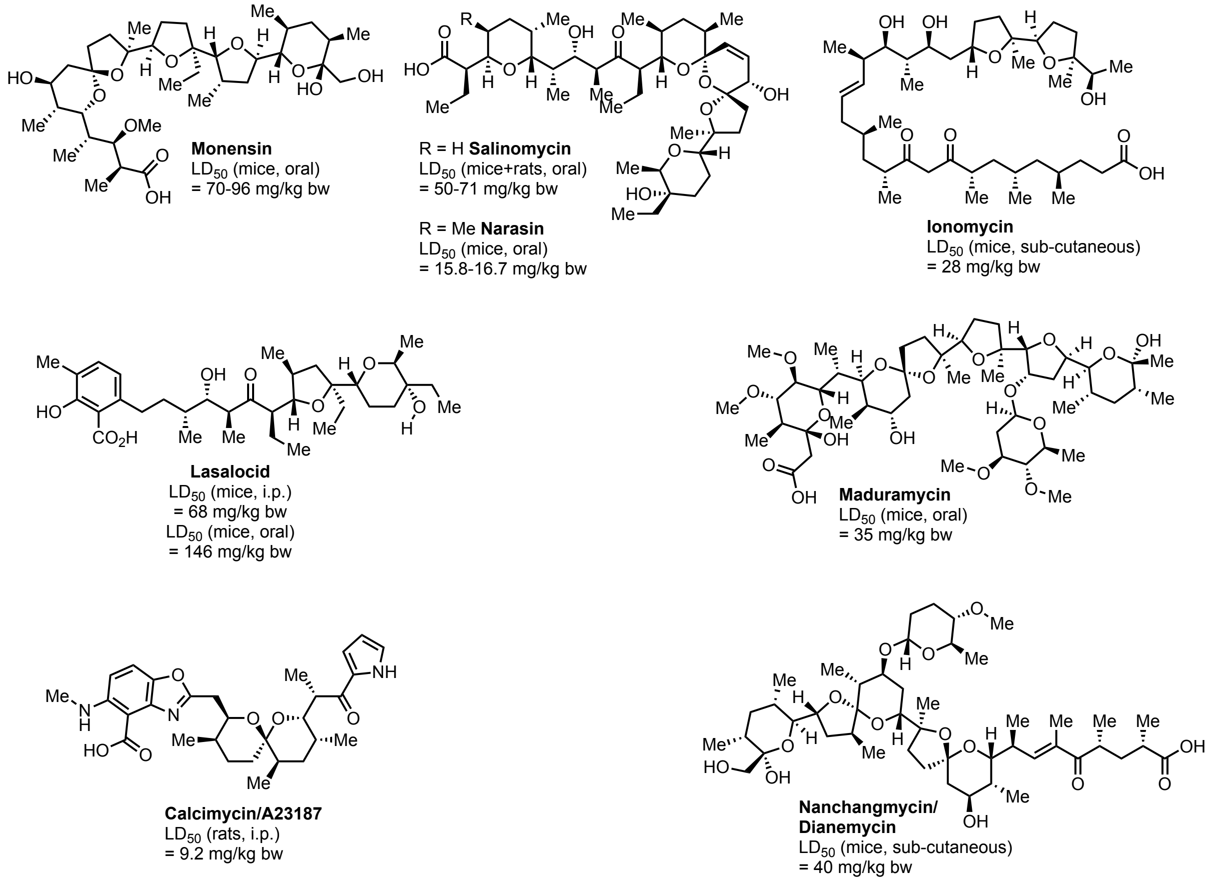


**Fig. S1.** Chemical structures of polyether ionophores used in this study and a compilation of selected, reported toxicity values (acute $LD_{50}$) in mice or rats. Additional data can be found in the parent resources: Monensin[1], salinomycin[2], narasin[3], Ionomycin[4], lasalocid[5], maduramycin[6], calcimycin[7], nanchangmycin[8].

---

**Table S1.** Biofilm eradication results in the clinical isolates. A biofilm in each strain was first grown for 24 hours and then treated for another 24 hours. The biofilms were allowed to recover for 72 hours, and the minimum biofilm eradication concentrations (MBEC) were determined and listed below in µg/mL. The assay was performed in five methicillin-sensitive strains and three MRSA strains. Clinical isolates 1, 2 and 5 are penicillin-resistant and methicillin-sensitive whereas clinical isolates 3 and 4 are both penicillin- and methicillin-sensitive. Clinical isolates 6-8 are methicillin-resistant. Wildtype *S. aureus* DSM 20231, and wildtype *S. aureus* MRSA USA 300 je2 were included for comparison.

| | Methicillin-sensitive *S. aureus* | | | | | | Methicillin-resistant *S. aureus* | | | |
|---|---|---|---|---|---|---|---|---|---|---|
| **Compound** | ***S. aureus* DSM 20231** | ***Clinical isolate 1*** | ***Clinical isolate 2*** | ***Clinical isolate 3*** | ***Clinical isolate 4*** | ***Clinical isolate 5*** | ***S. aureus* *MRSA* USA 300 je2** | ***Clinical isolate 6*** | ***Clinical isolate 7*** | ***Clinical isolate 8*** |
| Lasalocid | 64-128 | 128 | 128 | 128 | 128 | 128 | 128 | 128 | 128 | 128 |
| Salinomycin | ≥1024 | >1024 | >1024 | >1024 | >1024 | >1024 | 1024 | 1024 | 1024 | 1024 |
| Monensin | >128 | >128 | >128 | >128 | >128 | >128 | >128 | >128 | >128 | >128 |
| Calcimycin | 4-8 | 16 | 8 | 16 | 16 | 4 | 4 | 4 | 8 | 4 |
| Ionomycin | ≥1024 | >1024 | >1024 | >1024 | >1024 | >1024 | >1024 | >1024 | >1024 | >1024 |
| Narasin | 256-512 | 1024 | >1024 | >1024 | >1024 | 1024 | 256 | 256 | 256 | 256 |
| Maduramycin | ≥1024 | >1024 | >1024 | >1024 | >1024 | >1024 | >1024 | >1024 | >1024 | >1024 |
| Nanchangmycin | 32 | 64 | 64 | 128 | 64 | 64 | 128 | 128 | 8 | 8 |
| Vancomycin | 1024 | >1024 | 1024 | >1024 | >1024 | >1024 | >1024 | >1024 | >1024 | >1024 |

**Table S2.** List of genes with mutations found in nanchangmycin-resistant isolates. The sequencing data have been deposited to Genbank under Bioproject number PRJNA877072 and accession numbers: CP104478 (untreated reference), CP104477 (resistant mutant 1), CP104476 (resistant mutant 2), CP104475 (resistant mutant 3).

| Locus | Name | Predicted function |
|---|---|---|
| WP_000021864.1 | TrkH | Potassium uptake |
| WP_001161085.1 | MspA | membrane stabilizing |
| WP_000066900.1 | SarV | Transcriptional regulator |
| WP_000757569.1 | glycerophosphorodiesterase | Phosphodiester hydrolysis |
| WP_001151499.1 | ATP-dependent helicase recG | DNA repair |
| WP_001118443.1 | translation initiation factor IF-1 | Protein synthesis |

**Table S3.** Hits of the Nebraska Transposon Mutant Library screen, with corresponding MIC values of the relative gene-inactivated mutants against Lasalocid (Las), Salinomycin (Sal), Calcimycin (Cal), and Nanchangmycin (Nan).

| Locus | Name | Product | Las | Sal | Cal | Nan |
|---|---|---|---|---|---|---|
| MRSA wildtype | | | 4 | 2 | 0.0625 | 2 |
| SAUSA300_1357 | *aroC* | chorismate synthase | 0.25 | 0.125 | 0.0625 | 0.25 |
| SAUSA300_1615 | *hemB* | delta-aminolevulinic acid dehydratase | 0.5 | 0.25 | 0.0625 | 0.25 |
| SAUSA300_0844 | *ndh2* | NADH:ubiquinone reductase | 2 | 1 | 0.0625 | 1 |
| SAUSA300_0962 | *qoxB* | quinol oxidase, subunit I | 2 | 1 | 0.0625 | 1 |
| SAUSA300_0961 | *qoxC* | quinol oxidase, subunit III | 2 | 1 | 0.0625 | 1 |
| SAUSA300_0963 | *qoxA* | quinol oxidase, subunit II | 2 | 1 | 0.0625 | 1 |
| SAUSA300_1016 | *cyoE* | protoheme IX farnesyltransferase | 2 | 1 | 0.0625 | 1 |

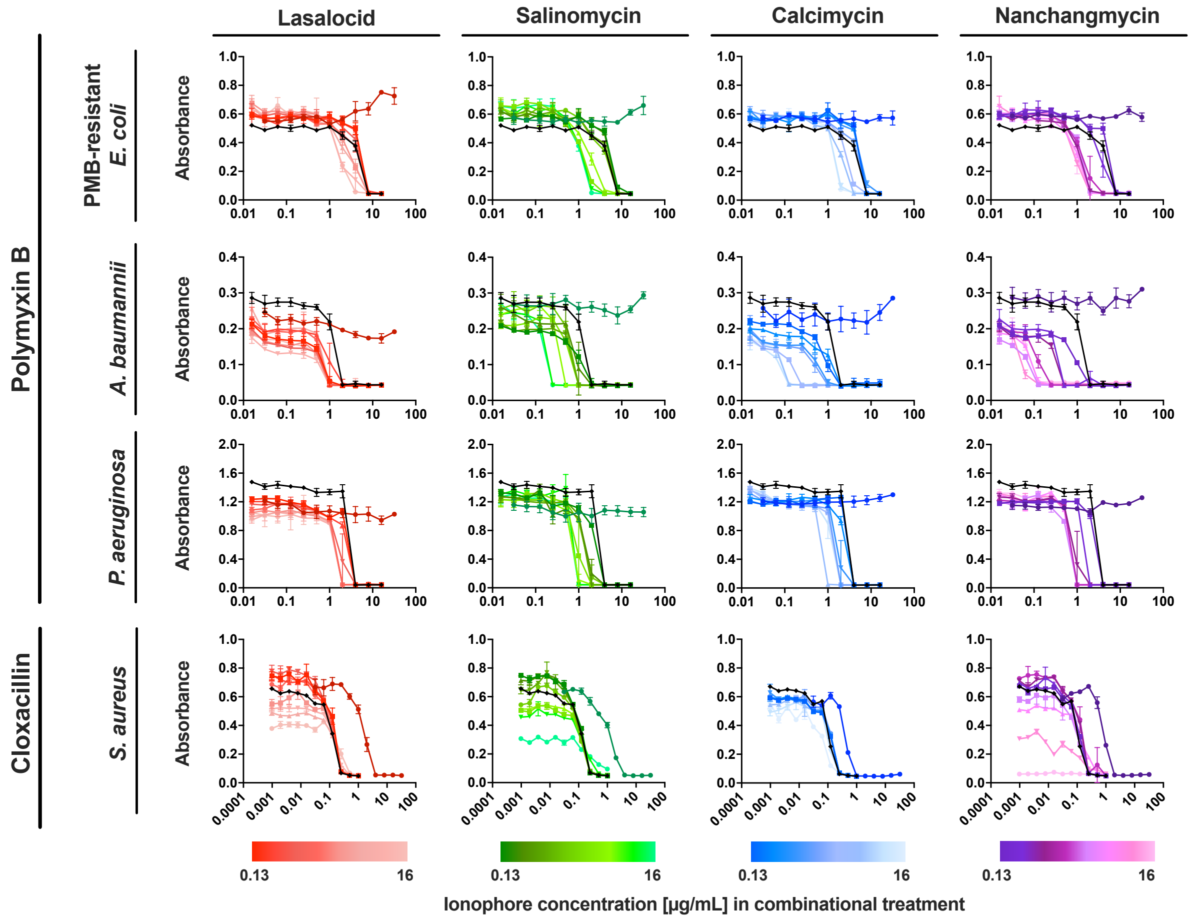


**Fig. S2.** Combinatorial treatment in a polymyxin-resistant *E. coli* (DH5α pGDP2 MCR-1), *A. baumannii* DSM 300007, *P. aeruginosa* DSM 19880, and *S. aureus* DSM 20231. The three gram-negative strains were treated with polymyxin B, and *S. aureus* was treated with cloxacillin, all in combination with four selected ionophores which were added as a fixed dosage (concentration range 0.13-16 µg/mL in gram-negative strains; 0.004-0.5xMIC in *S. aureus*). Polymyxin B, cloxacillin and the ionophores were tested as single treatments (PMB/cloxacillin: black; lasalocid: dark red; salinomycin: dark green; calcimycin: dark blue; nanchangmycin: dark purple) as positive and negative controls.
